# Monitoring *Candida albicans* Biofilm Formation by Impedance Using Passive RFID

**DOI:** 10.1101/2025.07.03.662933

**Authors:** V. Makarovaite, A. J. R. Hillier, S. J. Holder, C. W. Gourlay, J. C. Batchelor

**Affiliations:** Engineering Department, University of Kent, Canterbury, CT2 7NZ; Chemistry Department, University of Kent, CT2 7NZ; Bioscience Department, University of Kent, CT2 7NZ; Engineering Department, University of Kent, Canterbury, CT2 7NT

**Keywords:** RFID, Sensor, Yeast, Biofilm, Dielectric Properties

## Abstract

Medical implants are routinely colonized by microbial biofilms especially yeast biofilms such as those formed by *Candida albicans*. As medical implants move towards smart technology and IoT, it is more important than ever to understand the electrical properties of biofilm formation. Here we present an attempt at understanding how *C. albicans* biofilm formation effects sensor technology. We propose that as the biofilm matures, it becomes increasingly hydrophobic and undergoes a reduction in effective dielectric constant and ionic conductivity. This results in an impedance mismatch that disrupts capacitive coupling between the sensor and surrounding media, this effect that can be reliably detected. We show that a *C. albicans* biofilm changes its dielectric properties during maturation and, once mature, can be equated to a thin dielectric insulator. We demonstrate that these properties can be used to enable constant monitoring of a *C. albicans* biofilm using passive RFID, offering a new detection methodology that may be applied to surfaces known to be readily colonise.

## I. INTRODUCTION

**S**urface colonisation of implant material account for 20– 25% of primary complications [1], which with a 2025 projection for the global market in the medical implant sector exceeding US$100 billion with a US$1 billion increase on next generation implants alone [2, 3] could develop into a serious strain on the global healthcare system. Colonisation often leads to microorganism biofilm formation, where multiple (or singular) species adherence to a material surface causes extracellular matrix formation, which can even multiply to a 300 (or more)-micron thickness [4]. Bacterial and fungal organisms can be found within mixed biofilms that form on medical devices. The most common yeast species associated with biofilm formation and implant failures is *Candida albicans* [4]. *S. aureus* has been widely recognised as an implant coloniser, particularly on titanium surfaces [5], while *C. albicans* biofilms prefer colonisation of medical polymers [6]. Biofilm formation is clinically relevant as it allows microorganism(s) to become resistant to most drug interventions and the host immune system, further necessitating removal of the implant [6].

*C. albicans* biofilms are known to evade reactive oxygen species triggering and neutrophil killing within the host [7], to secrete aspartic proteases to avoid human complement attack [8], and employ other evasion methods that have been previously reviewed [4, 9]. *C. albicans* biofilms are known to cause over 100,000 deaths annually within the United States alone and are responsible for 40% of clinical blood infections and 15% of sepsis cases [10].

With the projected increase in next generation implants (or smart implants), it is becoming imperative to understand microbial biofilm formation in terms of electrical properties. Most of the biofilm growth detection systems rely on electrochemical impedance spectroscopy (EIS), which indirectly measures microbial growth by detecting changes in system conductivity [11]. While several studies have reported EIS-based impedance tracking for *S. aureus* biofilms [12–18], recent research has also demonstrated the feasibility of using impedance biosensors and microfluidic platforms for sensitive and real-time detection of *C. albicans* [25–27]. However, non-EIS impedance techniques (particularly low-cost, wireless, passive methods such as RFID) remain underexplored for fungal biofilms. This paper addresses this shortfall by demonstrating the application of passive RFID sensing to *C. albicans* biofilm formation and benchmarking the results against continuous EIS measurements.

In this study, we investigate the potential of passive RFID sensor systems to detect and monitor *Candida albicans* biofilm formation via impedance-related parameters. Our central hypothesis is that biofilm maturation alters the dielectric environment at the sensor interface, producing measurable changes in capacitance and impedance that can be captured through RFID-based sensing. To evaluate this, we utilise two complementary approaches: (1) real-time impedance tracking with the xCELLigence system to benchmark biofilm development, and (2) passive UHF RFID sensors to detect electrical property shifts associated with biofilm presence and growth. Additionally, we employ electromagnetic simulations to model the expected sensor responses based on biofilm thickness and composition. By correlating RFID sensor output with xCELLigence data and simulation results, we aim to validate the feasibility of using passive RFID technology for non-invasive biofilm monitoring in medical applications.

## II. METHODS

### A. Candida albicans Culture Conditions

The clinical isolate type strain SC5314 of *Candida albicans* was cultured on Yeast Extract Peptone Dextrose (YPD) agar (Sigma-Aldrich) with 2% glucose at 37^°^C for 48 hours. Then the cells were harvested and re-suspended in standard 2% glucose YPD liquid media (Sigma-Aldrich) followed by a 24-hour incubation at 37^°^C. This allowed for an overnight cell density growth of 10^7^ CFU/ml (haemocytometer counted) for the biofilm studies. Cell density was adjusted as needed.

### B. Biofilm Growth

For sterilisation, RFID wells were washed with a 70% ethanol solution and after drying, a 30-minute incubation with 50% Fetal Bovine Serum (FBS, Sigma-Aldrich) at 30^°^ C allowed for cell adherence. FBS helps coat the biofilm surface and allows for increased yeast adherence. After the FBS incubation, each well was washed with sterile PBS twice followed with an adhesion phase of 90 minute at 37^°^ C with 1 ml of *C. albicans* (10^7^ CFU/ml) in 10% glucose YPD media. After cell adhesion, each well was lightly washed twice with sterile PBS to remove non-adherent yeast cells and refilled with 1 ml of fresh YPD liquid media for a 48-hour incubation at 37^°^ C. The same process was used for xCELLigence detection of biofilm growth (detailed below) except for the sterilisation as all E-plate-units are shipped as sterile closed plates and they use 200 µl rather than 1 ml. xCELLigence standardized growth was then compared to two different RFID sensor designs for the ability to detect impedance changes with C. albicans biofilm formation.

### C. Assessment of C. albicans biofilm growth using the xCELLigence System

The real time cell analyzer (RTCA) xCELLigence (ACEA Bioscience Inc., San Diego, CA) detects growth of eukaryotic cells due to variations in impedance signals (stated as cell index (CI)) throughout the growth process upon the bottom of the gold-microelectrode E-plates [12]. This system has been previously utilised for biofilm growth detection of Staphylococcus aureus with repeatable results and a similar approach was taken for the presented *C. albicans* growth [13, 14]. The RTCA was placed in a 37^°^ C incubator with 5% CO2 without shaking during biofilm growth. The xCELLigence system was calibrated using 200 µl of fresh YPD media to have as the basal CI value of “0”. Once the basal value was recorded, the biofilm growth protocol was followed as described and the xCELLigence was run based on manufacturer guidelines [12]. To determine the *C. albicans* biofilm hydrophobicity, 2 µL or 6 µL of zymolyase (100T) was added to the mature biofilm at 24 hours equivalent to 1T and 3T, respectively. Growth of the biofilm correlates to change in detected impedance of the system. It should be noted that to determine the accuracy of the change in protocol for biofilm growth and adherence presented, *S. aureus* was utilized along with *C. albicans* and compared to [14]. *S. aureus* was utilized to determine the accuracy of biofilm growth using the RTCA as *S. aureus* xCELLigence growth has been previously published [13, 14] while no known xCELLigence biofilm growth publication exists for *C. albicans*.

### D. RFID Antenna Designs

Two main antenna designs were used to monitor biofilm growth (Fig. 1). The readily available RF-Micron RFM2100-AER [19] tag which utilises a Magnus-S3 chameleon IC able to adjust capacitance from 2pF to 3pF. It essentially acts as a sensing transducer able to compensate any reactance change by internal tuning via a switchable capacitor network output as a digital sensor code reading ranging from 32 to 0 for the 1pF change (Fig. 1 A). While the interdigit sensor tag utilises an Alien Higgs-3 Electronic Product Code Class 1 Gen 2 RFID integrated circuit (IC) (31-j216 Ω) [20] mounted on FR4 (Fig. 1 B). To assist in biofilm formation, both tags had a PDMS overlay of 20 µm following the protocol presented in [24] on the sensor regions and attachment of a square reservoir for the liquid growth media around the sensor region.

**Fig. 1.**
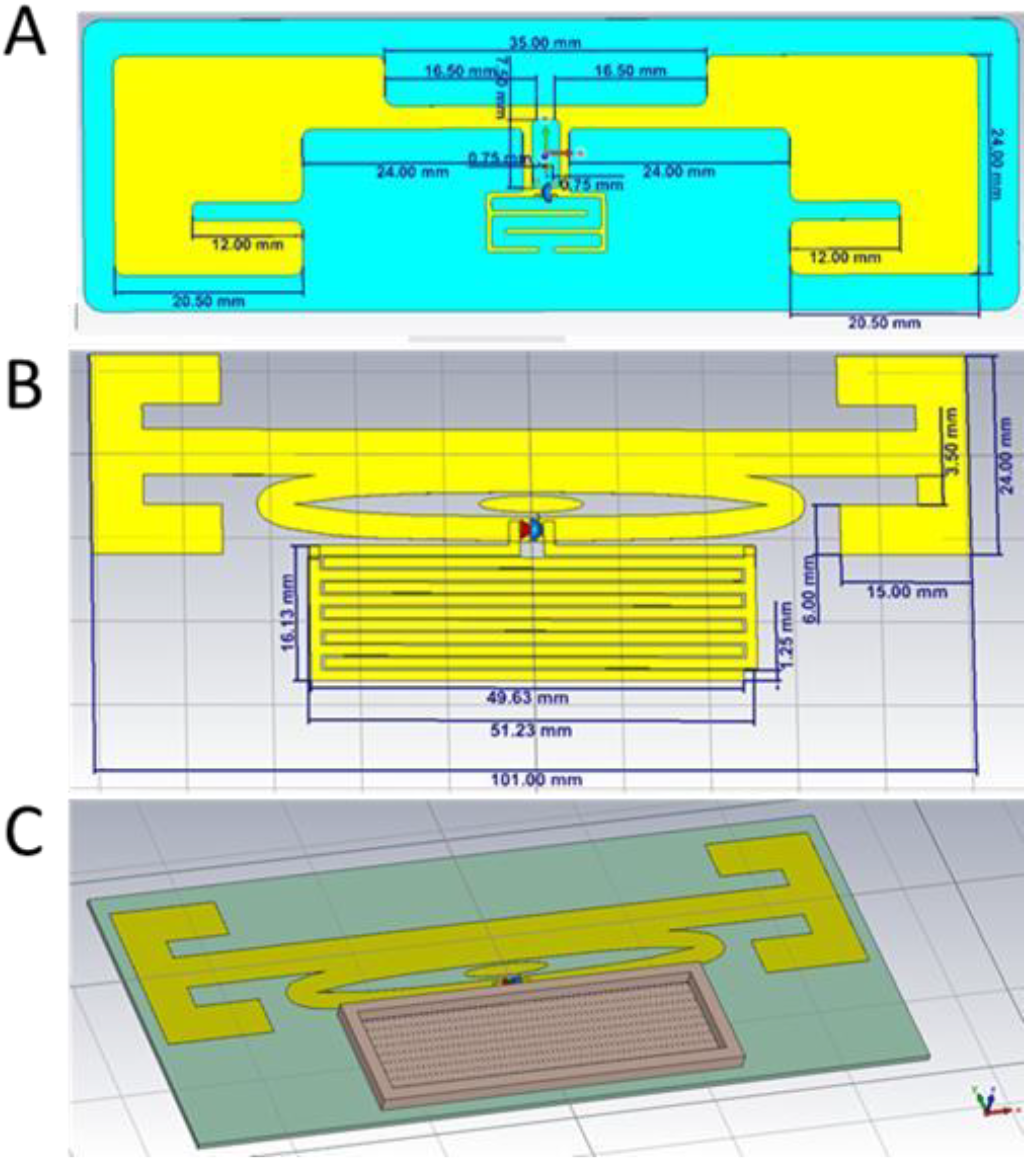
Antenna designs utilized within CST simulation: (A) RFMicron tag, (B) simple interdigit sensor for UHF antenna and (C) representation of the 5 mm height reservoir around the sensor region of (B) allowing for biofilm growth in liquid media

### E. RFID Measurements and Simulation

CST Microwave Studio ® was used to simulate *C. albicans* growth on the RFID design. The CST antenna models were adjusted with data obtained from the xCELLigence experiments (Fig. 2 and 3) to simulate a nonconductive layer of as a mature biofilm. This was also used to determine transponder chip and tag matching as well as predict the maximum theoretical read range. Antenna read range is the primary judge of RFID system performance and it determines the maximum distance that an RFID IC can be activated by passing the threshold for ‘wake-up’ power. The maximum read range was calculated by the formula:

**Fig. 2.**
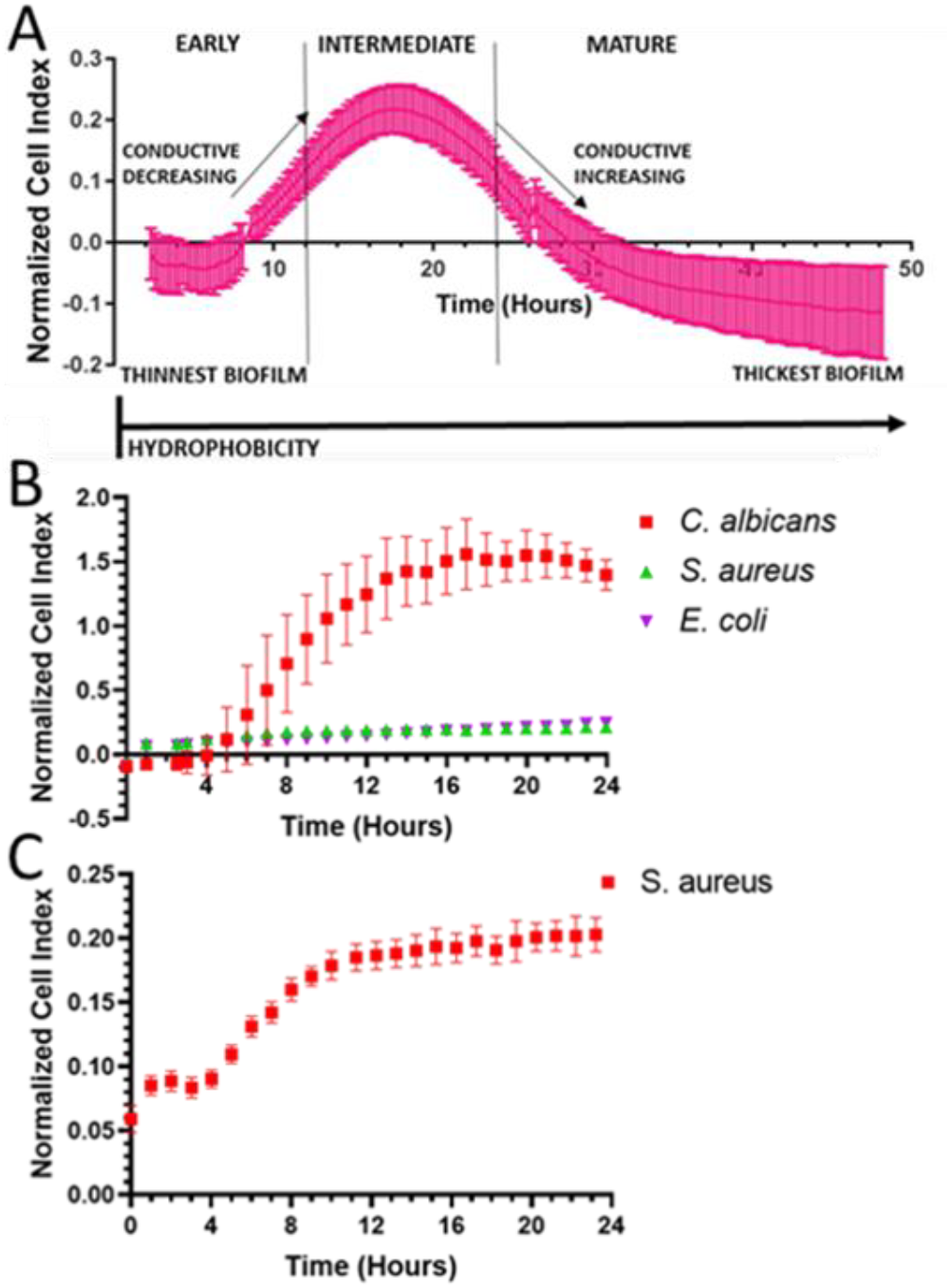
*C. albicans* biofilm growth with the impedance based xCELLigence system. (A) *C. albicans* 48-hour growth; the liquid media is represented as the 0 line on x-axis (N=3, n=18). The hours are broken down into early, intermediate, and mature biofilm hours where there is loss in conductivity which occurs starting at 18 hours. (B) 24-hour *C. albicans* growth in comparison to S. aureus (n=6). (C) *S. aureus* growth with a 0.10 cell index change (from minimum to maximum cell index values) (n=6). All data was normalized for evaporation and liquid media.

**Fig. 3.**
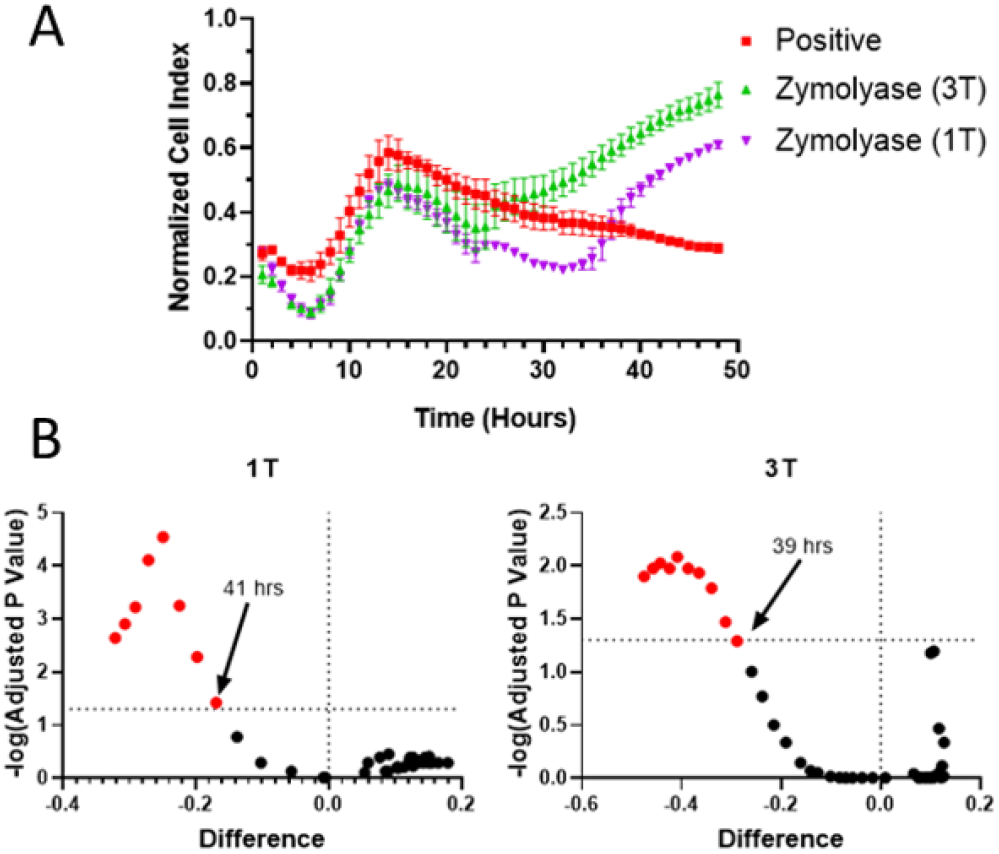
Zymolyase disruption of mature *C. albicans* biofilm growth within the xCELLigence system normalized against media control. A) *C. albicans* 48-hour growth with the addition of zymolyase at 24 hours; B) Volcanic plots showing first hour of significance (p-value < 0.05) with zymolyase (1T or 3T) addition at 24 hours during *C. albicans* biofilm growth in comparison to a positive control. Statistical significance determined using the Holm-Sidak method.

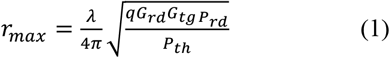

where Grd is the reader antenna gain, Gtg is the gain of the tag, q is impedance mismatch factor between the tag and IC, λ is the reader transmitted frequency, Prd is the reader transmitted power and Ptg is the received power by the tag [21]. The only flexibility in the system rests with changes to the Gtg or q of the system; all other aspects of the calculations are pre-set due to antenna reader limitations or manufacturer guidelines, and country laws dictating frequency use by RFID technology [21].

All RFID measurements were conducted using the Voyantic TagformancePro RFID measurement system in triplicates. The Voyantic system was set to sweep both the frequency (between 800MHz to 1000MHz) and power (from 0dBm to 30dBm) to determine the peak read range values. The *C. albicans* biofilm growth was measured after the removal of excess growth media using a pipette and air dried for 30min to reduce any stray conductivity not already present within the biofilm.

## III. RESULTS AND DISCUSSION

A correlation was observed between the results obtained with the xCELLigence system and the UHF RFID sensors, both of which suggest a link between mature biofilm hydrophobicity on the sensor surface and growth detection.

### A. Measurement of biofilm growth using xCELLigence

*S. aureus* growth was employed to test if the applied biofilm growth protocol would result in the same trends seen for biofilm forming wildtype *S. aureus* strains in [13] and [14]. This current protocol produced repeatable results comparable to [14] as the 0 to 6-hour (min) growth compared to the 18 to 24-hour (max) growth had a similar 0.12 CI difference (same as previously published).

With the xCELLigence RTCA, *C. albicans* growth showed a bell-curve appearance peaking at around 18 hours, where change in impedance plateaued before there was a decay of the impedance signal (Fig. 2 A and B). The reversal in impedance after 18h correlated with the formation of an intermediate to mature biofilm under these growth conditions [4-7]. This suggests that a key property of the biofilm during its maturation phase leads to a measurable change in sensor impedance. As shown in Figure 3A, the Cell Index (CI) declines after an initial peak (20hrs), indicating that the maturing biofilm increasingly insulates the electrode and disrupts the conductive pathway through the growth media.

As cells adhere and a biofilm begins to form on the electrode surface, they act as partial insulators, altering the local ionic environment and disrupting the baseline capacitive coupling between the electrode and the surrounding conductive media. This increases the magnitude of impedance measured by the xCELLigence system and results in a rise in Cell Index (CI) from its initial baseline value of zero, which corresponds to clean media with no biological coverage [12].

As the biofilm matures and secretes dense extracellular matrix material, it increasingly insulates the electrode, reducing ionic access and electrical coupling. At this stage, the impedance change plateaus or even decreases, causing a decline in CI, as the system becomes effectively decoupled from the surrounding conductive media. When zymolyase (Fig. 3A) enzymatically degrades the biofilm matrix, restoring ionic contact between the electrode and the bulk solution, impedance rises once more and CI values increase; indicating the recovery of capacitive and conductive pathways. This impedance-based signature of biofilm development and disruption is consistent with trends observed using UHF RFID sensor systems (Fig. 6), where comparable shifts in electrical response were detected.

After 24 hours of growth, the maturation of a *C. albicans* biofilm leads to deposition of a dense extracellular matrix and increased biofilm thickness [22]. Changes in the system’s electrical properties at this stage may be attributed to both the biofilm’s growing height and the insulating nature of its matrix.

Both factors likely reduce ionic mobility and limit the conductive interaction between the E-plate microelectrodes and the growth media. Supporting this, it is known that growth media is more conductive than fungal biomass [23], and that increasing yeast cell density reduces the overall conductance of the culture [23]. We suggest that as the biofilm matures, it increasingly acts like a dielectric insulator, progressively changing electric field density between the sensor and media.

The impedance behaviour of *C. albicans* biofilms can be described using a Cole–Cole-type model, where the dispersive dielectric properties of the maturing biofilm (comprising cells, extracellular matrix, and ionic gradients) result in frequency-dependent complex impedance. As the biofilm thickens, its effective capacitance decreases and the relaxation time distribution broadens, consistent with a shift in Cole–Cole parameters (e.g., increasing τ, decreasing permittivity, or α < 1) [28]. Such dynamics are observed as a drop in Cell Index in xCELLigence assays or loss of backscatter resonance in passive RFID tags.

It is plausible that the observed impedance reversal near the 20-hour mark in the xCELLigence system may, in part, result from the progressive depletion of conductive growth media, as nutrients are absorbed and metabolized by the proliferating *C. albicans* cells [4–7]. However, it is more likely that this impedance shift is driven by the deposition of extracellular matrix material as the biofilm matures. This matrix acts as an insulating layer that increasingly decouples the electrode from the conductive media, leading to a decline in Cell Index (CI). While [13] attributed changes in CI to the number of adhered cells (where greater cell attachment corresponds to higher CI), it is unlikely that a loss of surface contact explains the decline at 20 hours in our system. No shedding or detachment is typically reported at this growth phase, and biofilm dispersion in *C. albicans* is known to occur after approximately 48 hours [4-7]. Furthermore, it is important to note that the growth protocols used in [13] and [14] involved seeding biofilms into media already containing suspended (planktonic) bacteria, whereas our experiments were conducted using adhered *C. albicans* cells only. Thus, their impedance trends likely reflect a mixed response from both attached and suspended cells, unlike our system which isolates the impact of true surface-associated biofilm growth.

This can be further enhanced by [29] where the researchers determined that *C. albicans* mature growth can act as a capacitor (can store and release electrical charge) with increasing charge and discharge cycles as frequency is increased. This further proves our hypothesis because as *C. albicans* biofilms mature on sensor surfaces, their electrical behaviour begins to resemble that of an insulating dielectric layer. In early stages, the biofilm contains water and ions that support ionic conductivity, allowing measurable changes in impedance at lower frequencies. However, as the biofilm thickens and its extracellular matrix develops, it becomes increasingly dense and hydrophobic, limiting ionic mobility. Another explanation for the loss in signal may be loss of viability. However, *C. albicans* viability did not reduce over the period of biofilm growth examined as XTT assays performed (after the 48-hour growth in the xCELLigence E-plate) XTT near 2.2 ± 0.4 (Table 1).

**Table 1.** XTT ASSAY OF *C. albicans* BIOFILM FORMATION.

| Type | XTT assay<br>(Average) | Repeats |
| --- | --- | --- |
| Negative | $0.03 \pm 0.01$ | 24 |
| Positive | $2.2 \pm 0.4$ | 24 |
| 1T Zymolyase | $2.0 \pm 0.1$ | 5 |
| 3T Zymolyase | $2.3 \pm 0.2$ | 4 |

### B. Sensor Properties

Our simulations of the electric and magnetic fields show that there is an adequate response from the sensor designs to the biofilm growth (Fig. 4). The simulated surface current was able to produce 1732 A/m in current concentrated within the sensor region containing the biofilm at an 868 MHz frequency (Fig. 4 A). The electric field (Fig. 4 B) showed a maximum voltage produced of 879kV/m with the thickest *C. albicans* biofilm included in the simulation (89 µm). The magnetic density equated to 2312 A/ m concerted within the sensor region (Fig. 4 C). Similarly, the RFMicron design also had a good response for the magnetic field (383 A/m (Fig. 4 D)) and surface current (349 A/m (Fig. 4 E)). These suggest that the current design can produce good wave propagation.

**Fig. 4.**
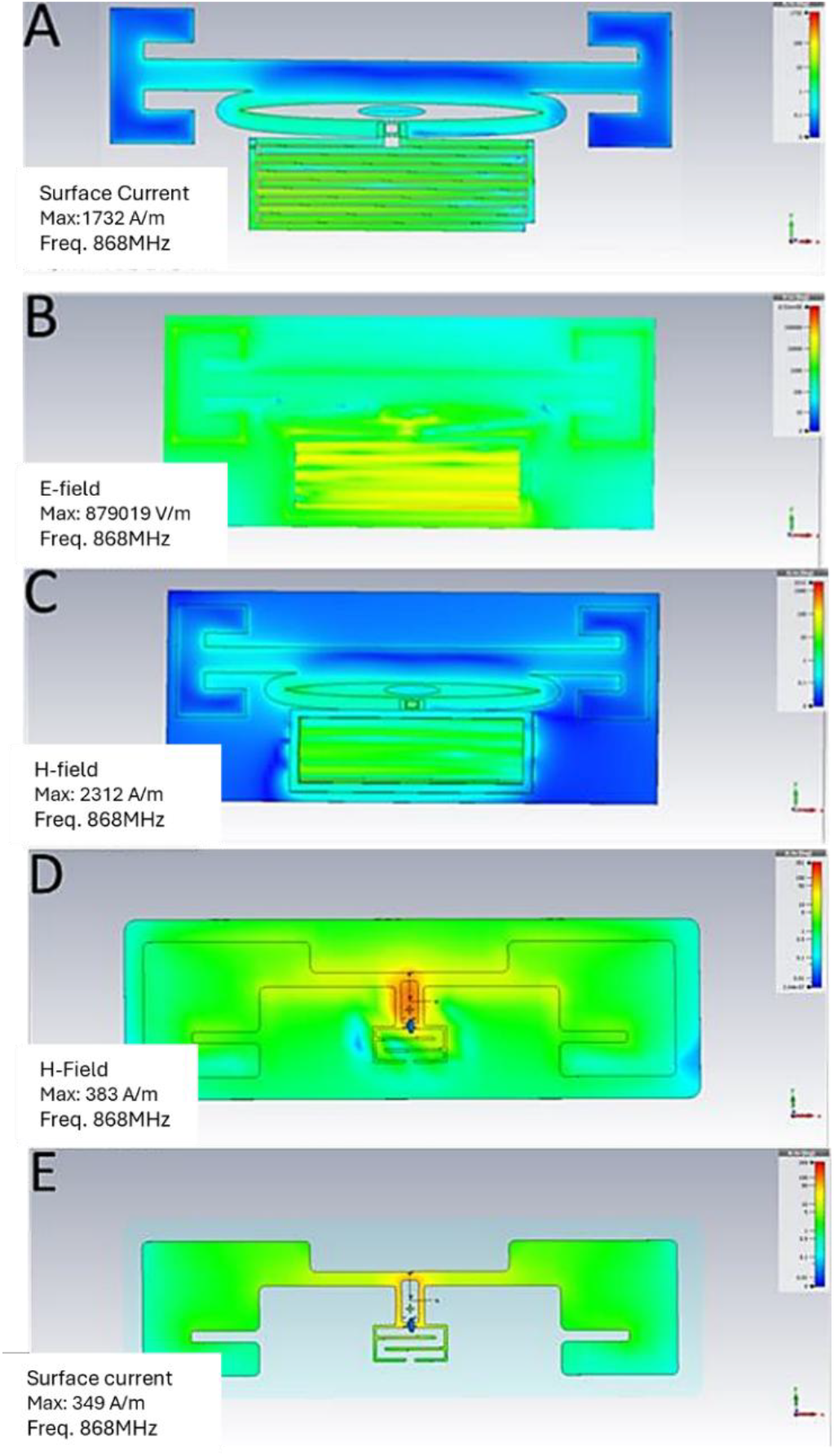
Simulated *C. albicans* biofilm growth effect on (A) surface current, (B) e-field and (C) h-field on the interdigit UHF sensor design. As well as the (D) h-field and (E) surface current for the RFMicron tag. Simulations exhibited are for the thickest biofilm growth studied (80 μm).

The simulated antenna directivity and realized gain (Fig 5 A and B, respectively). The directivity of the main lobe for the interdigit sensor UHF antenna was simulated as 1.89 dBi at a phi 90^°^cut, suggesting that this design has a primarily wide unidirectional appearance; however, it should be noted that the directivity flows in both the x and z axis with the majority of the read range reaching in the z-axis (Fig. 5 A). As for the realized gain of the antenna, the simulation calculated a low gain comparable to an electrically small antenna; though this can allow the antenna to be read from any location within 2 meters (x-axis) to 5 meters (z-axis). A highly directed antenna is not necessary for this study.

**Fig. 5.**
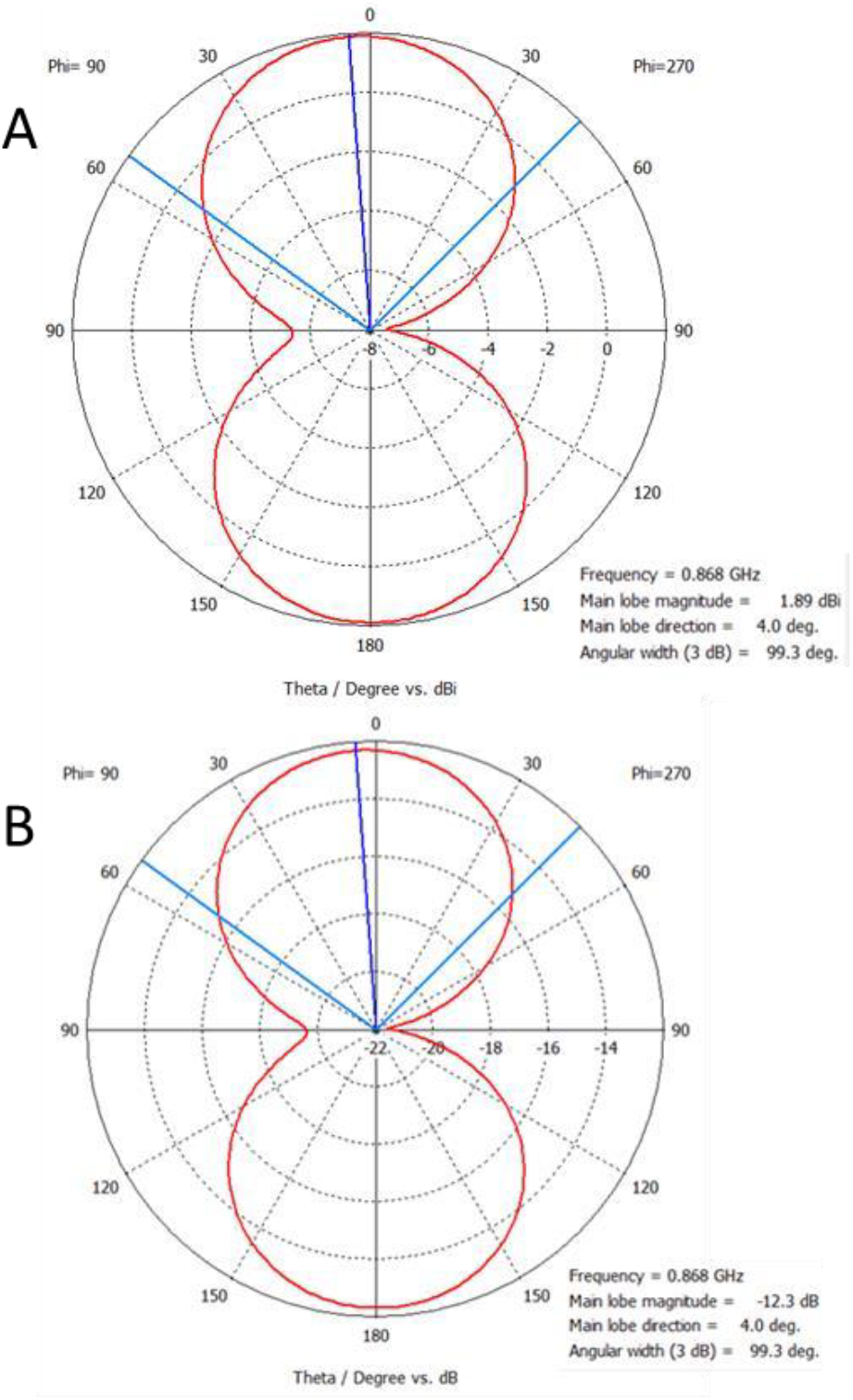
Simulated (A) directivity and (B) realized gain for *C. albicans* growth on the interdigit UHF sensor design. Both views were cut at phi 90^°.^

### C. *RFID Biofilm Growth of* C. albicans

To begin to investigate the cause of loss of impedance we simulated mature *C. albicans* growth on biosensors as an increase in a lossy insulator substance (simulated as increased layers of silicone (CST library) within this paper) based on the xCELLigence results. The thickness was adjusted from 0 µm to 80 µm as the majority of *C. albicans* biofilms remain within this thickness level [4-7]. It was seen that an increase in the biofilm thickness correlated with a change in capacitance exhibited as resonant frequency shifts with multiple resonance points in both simulation (Fig. 6A) and measured (Fig. 6B) for both the interdigital antenna design and the RFMicron design (Fig. 7A/B (simulated) and 7C (real)) which adjusted its internal capacitive tuning to account for the shift.

**Fig. 6.**
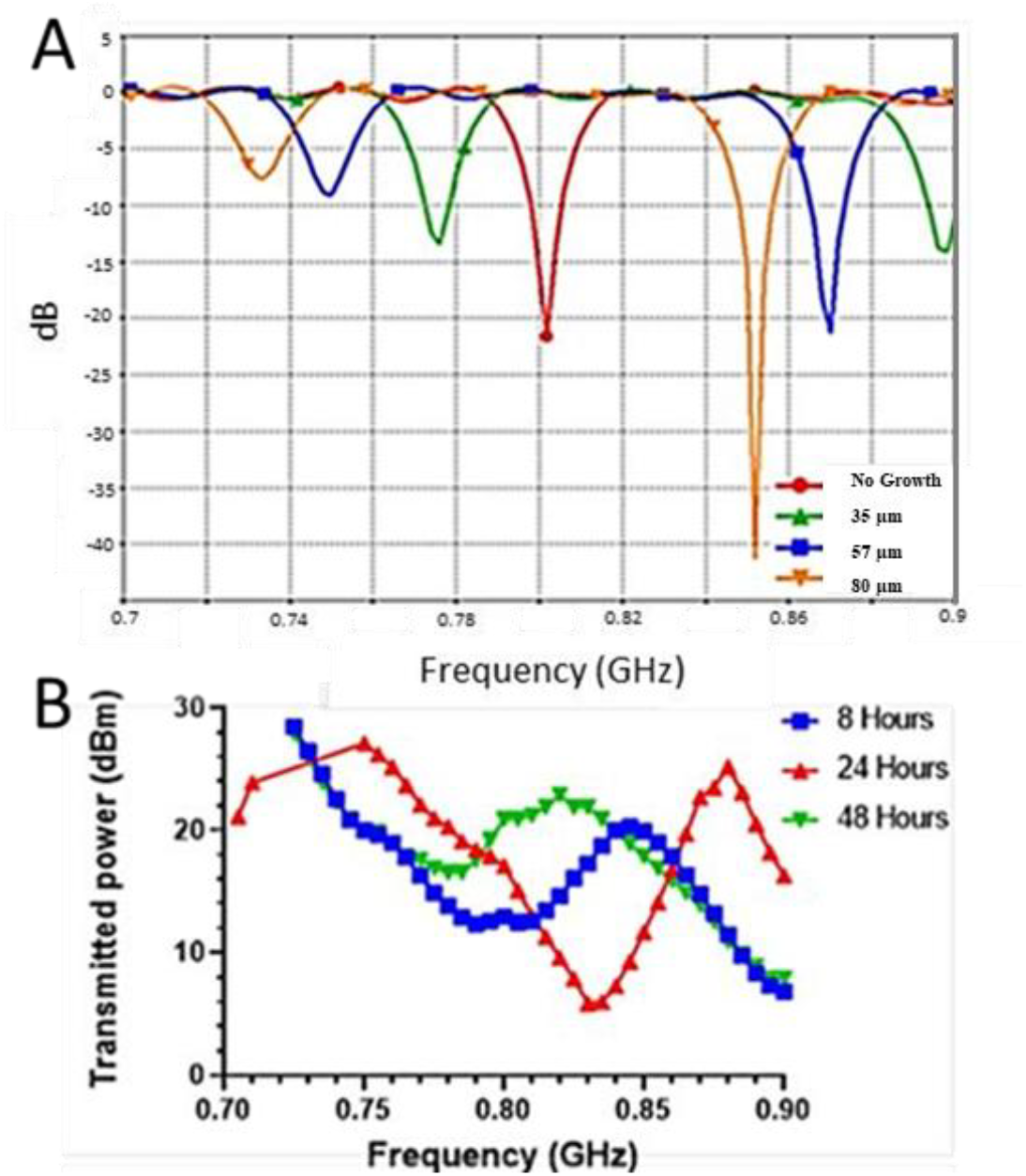
Interdigital RFID design. (A) S11 *C. albicans* growth simulation with increasing biofilm thickness on the interdigit UHF sensor design. (B) Measured *C. albicans* growth at 8, 24 and 48 hours with Voyantic TagformancePro; there was no noticeable change between clean media (0 hours) and 8 hours of growth. As biofilm grows in thickness or hours, there is a noticeable shift in frequency and more than one resonance points per thickness (n= 10).

**Fig. 7.**
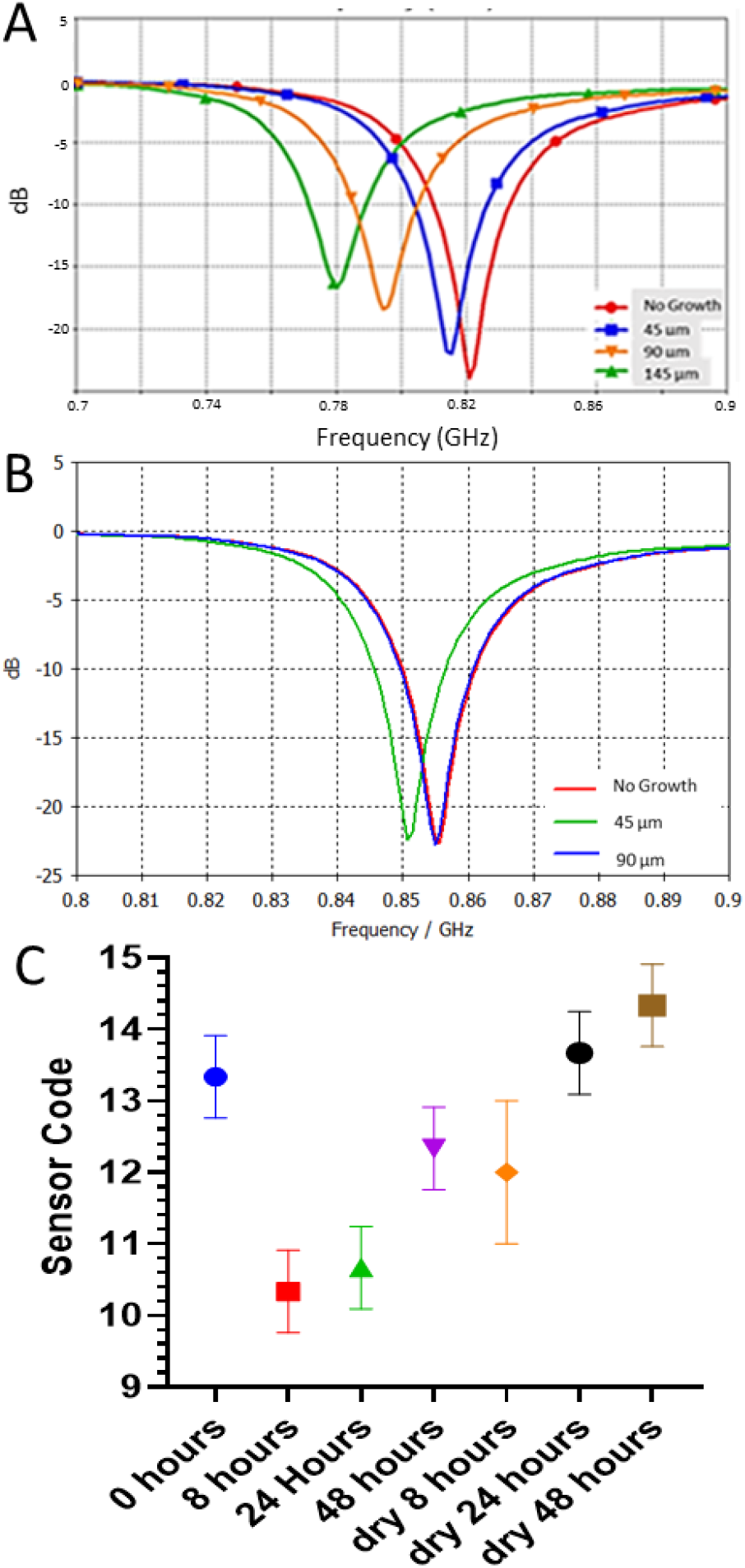
C. albicans growth effect on the RFMicron UHF sensor design. (A) S_11_ simulated increasing biofilm thickness without changing biofilm dielectric properties. (B) S_11_ simulated increasing biofilm thickness with changes in biofilm permittivity; no growth had permittivity equivalent to water, 45 μm decreased the permittivity to 47 and 90 μm had permittivity equivalent to 3 (2). (C) The RFMicron sensor tag measured detection of *C. albicans* biofilm growth.

We observed a large shift (near 100 MHz) from a thickness with 0 growth to 35 µm; at the second resonance point, there was a decrease in resonant frequency of about 20 MHz (Fig. 6A). This trend continued up to 80 µm thickness. This dual resonance was also seen with the ‘no growth’ thickness which had the second resonance at 925 MHz (not shown). Depending on resonance frequency used per thickness, this RFID sensor would produce a bell-curve (regarding biofilm thickness) that is comparable to that obtained using the xCELLigence system (Fig. 4 A). If looking at capacitance (within the 0-hour growth resonant frequency bandwidth), our simulations suggested that an increase in biofilm thickness of 35 µm would result in a decreased capacitance and a near 30Ω resistance loss (Table 2). With further biofilm increase in thickness, our simulations suggested that the capacitance would increase while there was a continued decrease in resistance.

**Table 2.** SIMULATED INTERDIGITATED ELECTRODE IMPEDANCE WITH INCREASING *C. albicans* BIOFILM THICKNESS AT 868 MHz.

| Thickness ($\mu\text{m}$) | Capacitance | Resistance |
| --- | --- | --- |
| No growth | $1.9\text{e-}10$ | 34.9 |
| 35 | $1.2\text{e-}12$ | 4.5 |
| 57 | $1.8\text{e-}12$ | 0 |
| 80 | $2.0\text{e-}12$ | 1.2 |

As Table 2 values were read at a constant 868 MHz to determine the effect of thickness on the sensor design, this does not consider the real changes that would occur when measuring the sensor. To achieve peak resonance matching, a shift in frequency to compensate for the IC impedance matching would occur. This correlates with the trends seen in all sensors (Fig. 8) as the ‘no growth’ and 48-hour growth had the highest capacitance simulated. It should be noted that in RF impedance calculations for antenna and IC matching, frequency and capacitance tend to have an inverse relationship; as capacitance increases in the system, the resonant frequency must decrease to stabilize the system at a certain impedance.

**Fig. 8.**
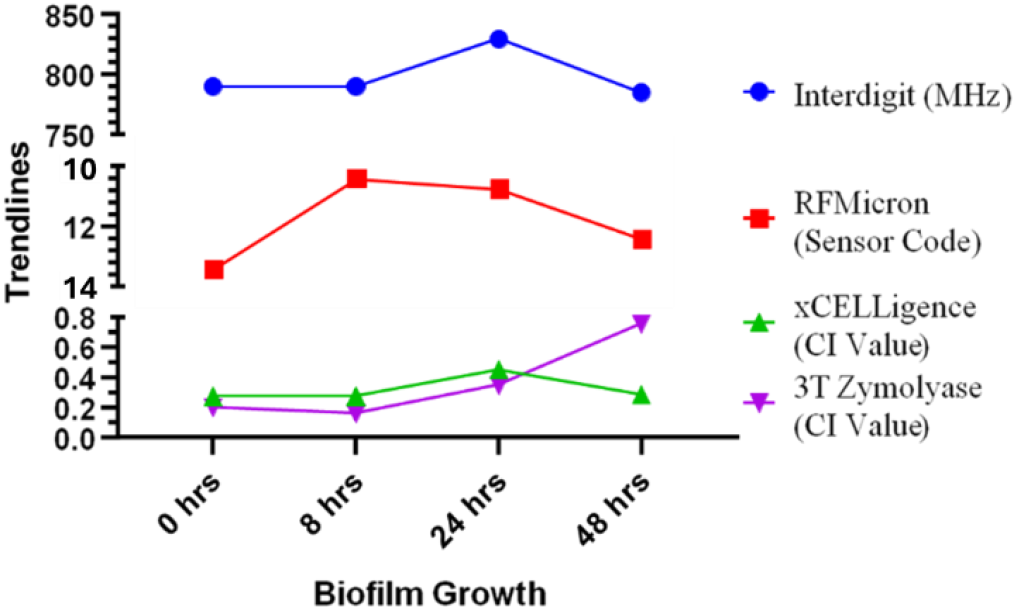
Measured data for all methods compiled showing changes corresponding to *C. albicans* biofilm growth. All methods demonstrated a distinct shift in measured signal from 0 to 24 hours, consistent with early biofilm growth and increased impedance. From 24 to 48 hours, a reversal or plateau in signal was observed, with 48-hour values closely resembling baseline (0 or 8 hour) readings. This pattern is consistent with sensor decoupling due to mature biofilm formation. The only deviation from this trend occurred in the zymolyase-treated samples, where continued signal increase between 24 and 48 hours reflected enzymatic biofilm disruption and re-establishment of capacitive coupling, evidenced as an increase in Cell Index.

When looking at the measured values using the Voyantic TagformancePro antenna reader system (Fig. 6B), we observed a similar shift in resonant frequency with increase in biofilm growth, where the biofilm acts as a progressively more insulating layer with reduced dielectric constant. This trend was similarly reflected in the RFMicron sensor (Fig. 7C), where the internal IC capacitance (reported as sensor code) changed in response to biofilm accumulation. Simulations of the RFMicron sensor (Fig. 7A), where increased insulating layer thickness was modelled using a uniform low-dielectric material, showed a comparable frequency shift consistent with a reduction in capacitive coupling. In Fig. 7B, simulation conditions were tailored to mirror the xCELLigence system response: the “No Growth” case represented highly conductive liquid media (highest dielectric constant); the intermediate biofilm thickness (45 µm) included partial water channels [4–7], maintaining some conductivity; whereas the mature biofilm (90 µm) was treated as a dense, low-permittivity layer, resembling a dehydrated insulator (lowest dielectric constant). As biofilm thickness increased, the system exhibited a bell-shaped impedance response, indicating that mature *C. albicans* biofilms functionally behave as low-dielectric, insulating layers, reducing effective field coupling and sensor resonance; this trend that matches observations across both RFID and xCELLigence platforms.

The RFMicron tag, there was a significant difference between the 0 hour (no growth), 8 hours and 48-hour growth when the samples were air dried (Fig. 7 C). The same bell-curve response was noted as seen with the xCELLigence; the capacitance decreased from 0 hours to 24 hours before it reversed within 24 to 48 hours (7 C). When the samples where fully dried with ethanol (Fig. 8), there was less significant variation between the dry growth hours and the 0-hour growth control, yet, the change in sensor code corresponded to what was expected when there is only an increase in a polymer layer on top of the RFMicron sensor [24]. This trend correlates exactly to what was seen in measured values of the xCELLigence (Fig. 2A).

At the set frequency of 868 MHz, there is a noticeable drop in resistance (Table 2) not because the material becomes more conductive, but because the system compensates for the increased impedance mismatch by adjusting capacitance and/or shifting the resonant frequency which can be seen in the simulated designs (Fig. 6A and 7A).

In sensing methods where the frequency is fixed, this compensation primarily occurs through changes in capacitance instead. This behaviour is consistent with observations from both the xCELLigence system and RFMicron RFID sensors. Moreover, the zymolyase-treated data (Fig. 7C) suggests that the presence of fluid within the biofilm matrix strongly influences sensor response. When this fluid is removed, the dry biofilm behaves more like a uniform dielectric, and capacitance displays a linear relationship with thickness; matching predictions from simulations (Fig. 7A). Collectively, these results suggest that as biofilm formation progresses, increasing matrix deposition and cell density render the surface more hydrophobic and insulating.

## IV. CONCLUSION

*C. albicans* biofilm growth detection is possible utilising both RFID UHF antenna sensors and the purchased xCELLigence equipment. Both system types were able to detect biofilm formation at intermediate and mature stages. The xCELLigence with the ability to do continuous (30-minute intervals) monitoring of growth was able to detect changes before the 24-hour mark. In theory, both the interdigit sensor and RFMicron UHF designs should, also, be able to detect early growth however some system tuning would be necessary especially if detection within growth media was necessary rather than taken as a snapshot monitoring. RFID offers an inexpensive high-performance alternative that performs as well as established technology.

